# dSreg: A bayesian model to integrate changes in splicing and RNA binding protein activity

**DOI:** 10.1101/595751

**Authors:** Carlos Martí-Gómez, Enrique Lara-Pezzi, Fátima Sánchez-Cabo

## Abstract

Alternative splicing (AS) is an important mechanism in the generation of transcript diversity across mammals. AS patterns are dynamically regulated during development and in response to environmental changes. Defects or perturbations in its regulation may lead to cancer or neurological disorders, among other pathological conditions. The regulatory mechanisms controlling AS in a given biological context are typically inferred using a two step-framework: differential AS analysis followed by enrichment methods. These strategies require setting rather arbitrary thresholds and are prone to error propagation along the analysis. To overcome these limitations, we propose dSreg, a Bayesian model that integrates RNAseq with data from regulatory features, e.g. binding sites of RNA binding proteins (RBPs). dSreg identifies the key underlying regulators controlling AS changes and quantifies their activity while simultaneously estimating the changes in exon inclusion rates. dSreg increased both the sensitivity and the specificity of the identified alternative splicing changes in simulated data, even at low read coverage. dSreg also showed improved performance when analyzing a collection of knock-down RBPs experiments from ENCODE, as opposed to traditional enrichment methods such as Over-representation Analysis (ORA) and Gene Set Enrichment Analysis (GSEA). dSreg opens the possibility to integrate a large amount of readily available RNA-seq datasets at low coverage for AS analysis and allows more cost-effective RNA-seq experiments. dSreg was implemented in python using stan and is freely available to the community at https://bitbucket.org/cmartiga/dsreg.

## Introduction

Eukaryotic genes are generally constituted by exons and introns (25). Alternative mRNAs may be generated from the same gene by inclusion or skipping of a particular exon in the mature transcript, in a process known as alternative splicing (AS) (19, 36). There is evidence of AS for most mammalian genes (35, 38) and of widespread changes of AS patterns throughout brain and heart development (4, 5, 15, 18, 23, 42, 43, 56). Defects in mRNA processing of some specific genes often lead to disease (5, 26, 27) and have been associated with complex neurological disorders, such as autistic syndrome (23, 28, 42, 55), and cancer (12, 49). Therefore, understanding the regulatory mechanisms underlying physiologic and pathological changes in AS patterns is crucial, not only to understand RNA biology, but also to identify potential therapeutic targets with a more general effect in complex diseases.

A two step work-flow is generally applied to identify the regulatory mechanisms underlying the changes in AS (see Figure 1 for a schematic representation). First, AS changes must be identified. For this, short reads from RNA sequencing are typically mapped using splice junctions (SJ) aware aligners such as STAR or Hisat2 (13, 40). Alternative mRNA processing can be studied at two different levels: 1) transcript quantification level, which can be based on a prior alignment (40, 52), or can be directly estimated from fast pseudoalignment methods (9, 39); and 2) event level quantification, as performed by popular tools such as MISO, MATS, vast-tools, DEXseq or SUPPA (1, 3, 23, 24, 46, 53). Recent toolsshowed improved performance for the estimation of AS changes by using information about exon features and perturbation experiments (21, 58). Since regulation is expected to take place locally, the event level analysis is the preferred approach to study the regulatory mechanisms underlying changes in AS profiles. Once AS events have been identified and quantified, different statistical approaches can be applied to determine AS changes between biological conditions, being a Generalized Linear Model (GLM) with binomial likelihood the most natural parametric approach (46).

**Fig. 1.**
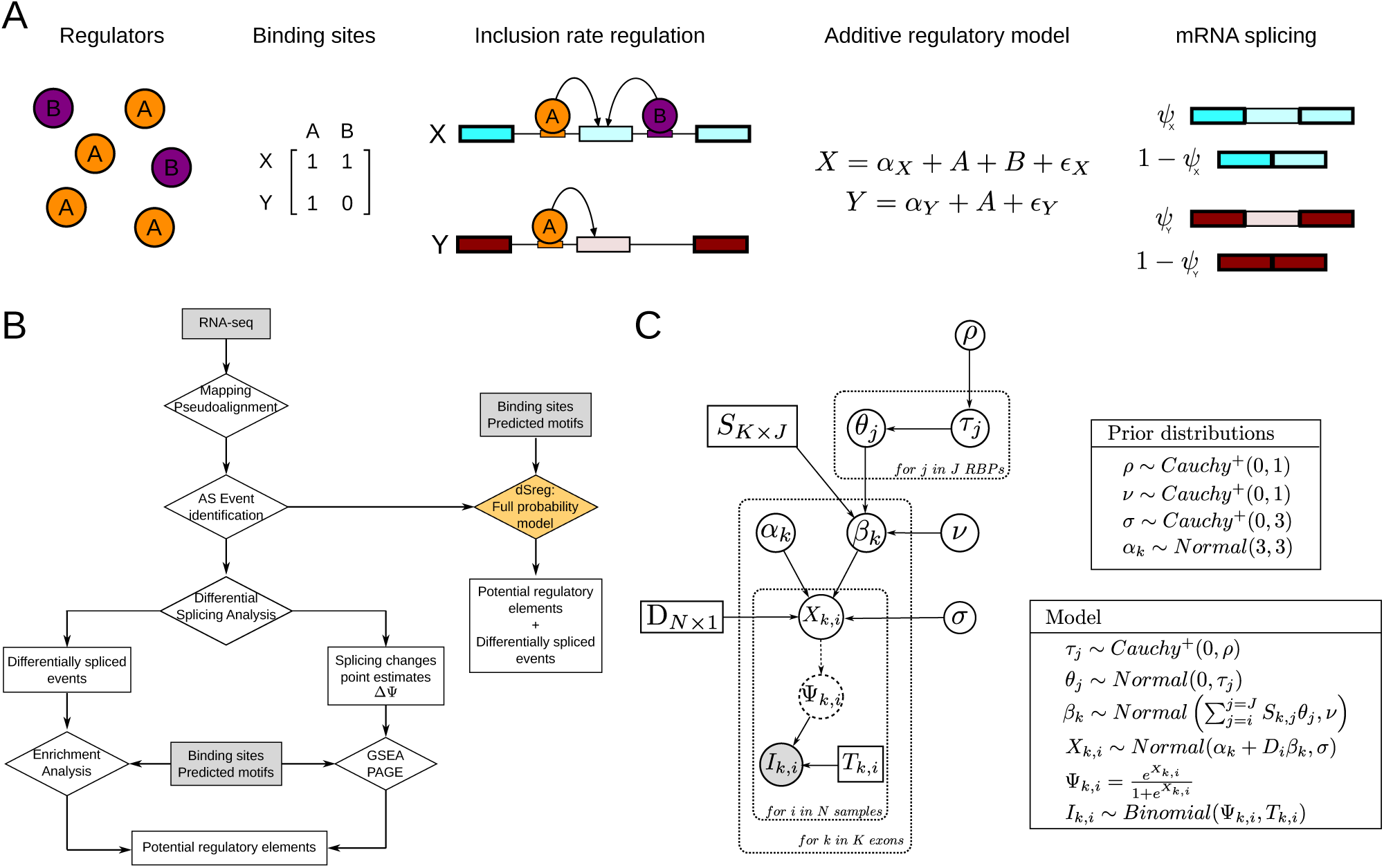
General and proposed work-flows for AS regulation analysis. **A** Schematic representation of idealized model for the regulation of splicing rates by RBPs through direct binding to their binding motifs in the pre-mRNA **B**. Diagram representing the different steps required for a classical analysis of regulation of alternative splicing using RNA-seq data and the proposed model in dSreg. **C**. Directed Acyclic Graph (DAG) representing the full probabilistic model integrating both differential AS analysis with binding sites presence and changes in the activity of RBPs.

The second step aims to identify regulatory features, such as RBPs motifs, associated with AS changes. Such features often include nucleotide hexamers, predicted motifs, experimentally determined or predicted binding sites (14, 17, 44, 57). Over-representation Analysis (ORA) enables the discovery of features co-ocurring with significant AS changes more often than expected by chance. Therefore, a sufficiently large set of significantly changed events is required to reach enough power to detect enrichment of regulatory features. ORA requires the categorization of splicing changes into different groups e.g. included or skipped, ignoring quantitative information about AS changes. To make use of quantitative information in the enrichment procedure, several approaches have been developed, including the widely known Gene Set Enrichment Analysis (GSEA) (48, 50). Although these tools were designed for functional analysis, they have been used to perform enrichment of known targets of regulatory elements (45, 53). However, the inherently noisier nature of the estimation of differences in AS compared to those of differential gene expression may limit the applicability of GSEA-like methods. Moreover, an additional limitation affecting both ORA and GSEA approaches lies on the high number of different features or binding sites and on the potential co-linearities among them that should be considered, resulting otherwise in a high false positive rate derived from confounding effects.

In this work, we used simulated data to study, for the first time, the performance and limitations of the classical enrichment approaches (ORA and GSEA) for the detection of regulatory elements driving AS changes. To tackle some of these limitations, we developed dSreg, a probabilistic model integrating differential splicing and regulation analyses. dSreg models latent changes in inclusion rates as a linear combination of the regulatory effects of the RBPs binding to relevant regions of the pre-mRNA for every identified AS event. Moreover, we used a hierarchical shrinkage prior distribution to model the changes in the activity of RBPs to formalize the assumption that only a few RBPs would show changes in their activity. dSreg was applied to simulated and real data, including data from systematic RBPs knock-down experiments, to assess its performance against ORA and GSEA. Finally, we applied dsReg to a real RNA-seq dataset obtained from a cardiomyocyte differentiation experiment, for which a limited number of AS regulators might be assumed.

## Methods

### dSreg: a mechanistic probability model for differential splicing

dSreg models the AS changes between two different conditions, *a* and *b*, as a function of changes in the activity of a few of the existing RBPs acting through their known binding sites. Given *K* AS events detected across *N* samples, we observe *I*_*k,i*_ reads supporting exon inclusion out of a total of *T*_*k,i*_ reads mapping to the *k*^*th*^ exon skipping event in sample *i*, which depends on the unknown probability of inclusion Ψ_*k,i*_. The conditional probability of observing *I*_*k,i*_ reads given *T*_*k,i*_ and Ψ_*k,i*_ is given by the binomial distribution.

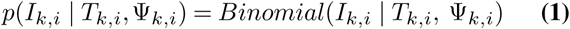

Ψ_*k,i*_ is therefore different for each sample *i*, but depends on the condition or group to which it belongs. Since probabilities are bound between 0 and 1, to model this dependency, we take the logit transformation *X*_*k,i*_,

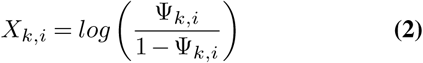

We assume that *X*_*k,i*_ is drawn from a normal distribution with a common standard deviation *σ*_*k*_ and different means per condition: *α*_*k*_ for condition *a*; and *α*_*k*_ + *β*_*k*_ for condition *b*, such that *β*_*k*_ represents the difference between the two conditions. For simplicity, we assume here that the standard deviation is the same across all *K* AS events (*σ*_*k*_ = *σ*).

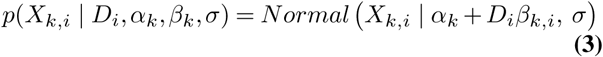

where *D*_*i*_ is a constant that takes the value 1 when the sample belongs to condition *b*, and 0 when it belongs to condition *a*:

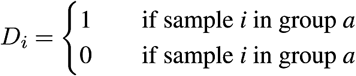

So far, this model is a simple logistic regression for each event with the only assumption that the sample variance is common across events and conditions. However, the changes in the probability of inclusion of exon *k* between two conditions, indirectly modeled by *β*_*k*_, should depend on the change in the activity *θ*_*j*_ of a particular regulatory RBP *j* and on whether it can bind to exon *k*. The binding information is encoded in a matrix **S**_*K*×*J*_, with value 1 whenever the RBP *j* binds to the exon *k* and 0 otherwise. Position dependent effects can be easily included by considering RBP *j* binding to different relative locations as different and independent RBPs. At the same time, the matrix **S** could also contain continuous values such as the probabilities of binding, affinities or scores given by Position Weighted Matrices (PWMs) (44) or any other predictive tool (2, 33).

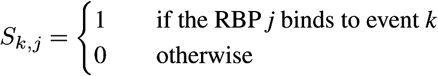

Now we can model *β*_*k*_, the change in the logit-transformed inclusion rate of exon *k*, as a normal distribution centered at a linear combination of regulatory effects 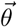 and 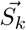 (the binding profile of exon *k*) with certain standard deviation *ν*. Adding variance *ν* to the distribution of *β*_*k*_ allows the existence of some changes in AS not necessarily explained by the regulatory features included in the model.

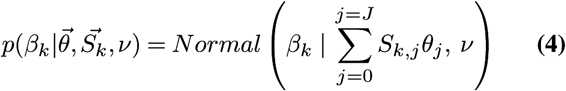

In this type of exploratory analysis, large numbers of regulatory proteins are usually tested. However, we expect that AS changes are driven by only a few RBPs. We formalize this prior belief setting a horseshoe prior for the change in the activity of regulator *j θ*_*j*_(11). The horseshoe prior, a member of the family of hierarchical shrinkage priors, specifies a normal prior for *θ*_*j*_ with mean 0 and a standard deviation *τ*_*j*_, where *τ*_*j*_ is not a fixed value, but drawn from a common half Cauchy distribution with mean 0 and *ρ* standard deviation. *τ*_*j*_ represents a local shrinkage parameter, as it only affects protein *j*, whereas *ρ* can be understood as a global shrinkage parameter. We further set a half Cauchy prior in *ρ* with mean 0 and standard deviation 1 as recommended (11). Note that this prior can be adapted according to the expected number of non-zero parameters (41).

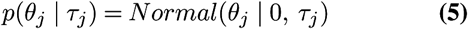

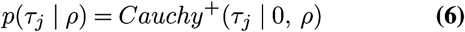

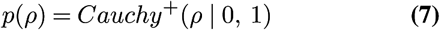

Finally, we need to specify prior distributions for the remaining parameters *α*_*k*_ and *σ*. Since we expect most of the exons to be included most of the times (Ψ ∼ 1) and *α*_*k*_ is the logit transformation of the inclusion rate in condition *a*, we set a normal prior centered at 3 (which reflects an expected Ψ = 0.95), with standard deviation 3 for each exon *k* to enable some deviation from this expectation. Moreover, as we expect little variation among samples, we set a half Cauchy prior distribution with 0 mean and standard deviation 1 on *σ*.

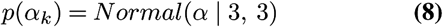

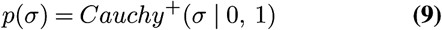

The joint posterior probability of the parameters Θ given the data (*I*) is proportional to the joint probability distribution of the data and Θ, since the marginal probability of obtaining the data *p*(**I**) is constant for any Θ.

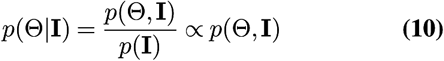

Using the conditional probabilities and prior distributions that we have defined for each variable, we can calculate this joint probability distribution applying the chain rule.

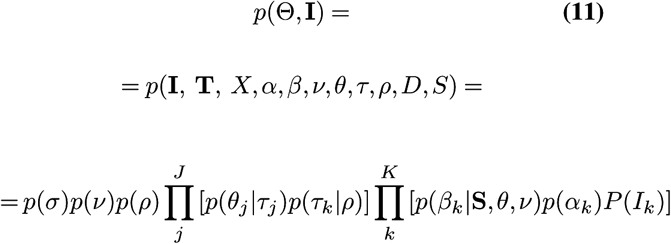

where,

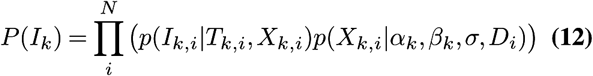

Once the full posterior distribution is completely specified, it can be explored using Markov Chain Monte Carlo (MCMC) algorithms. We implemented this model in stan (10), using a non-centered parametrization whenever possible to alleviate sampling difficulties from hierarchical models (7). The full model is represented as a Directed Acyclic Graph (DAG) to show dependencies among parameters in Fig. 1B.

### Data simulation

Data can be simulated by setting fixed values on the parent nodes of the DAG (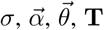 and **S**) representing the probabilistic model (Fig. 1C) and drawing samples from the corresponding distributions for each parameter. We simulated 20 datasets for each initial set of conditions, all with K=2000 events, 3 samples per condition (N=6) and J=50 potential regulatory elements with correlated binding profiles, of which only 5 showed non-zero effects on splicing changes between the two conditions.

To simulate realistic values of inclusion rates for the condition *a* (Ψ_*k,a*_) across the K=2000 exons, we assumed that 20% of the exons are alternative, with inclusion rates following a uniform distribution between 0 and 1; and 80% are consitutive, with inclusion rates drawn from a Beta(10, 1), to promote generally high inclusion rates.

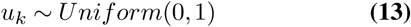

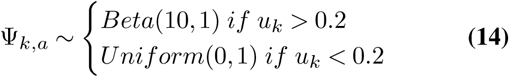

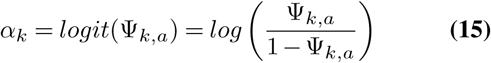

We aimed to simulate matrices of correlated binding profiles to take into account that certain groups of RBPs often bind to similar regions in the exons. To do so, we first simulated a covariance matrix Σ of size J sampling from an inverse Wishart distribution,

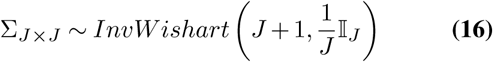

and used it to simulate K samples from a multivariate normal distribution using a mean of −2.5. This value represents an expected 7.5% of events bound by a particular RBP.

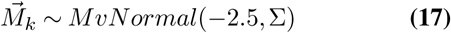

Then, we took the inverse logit to transform **M** matrix into the probability matrix **P** and use these probabilities to simulate binary binding profiles across exons (**S**_*K*×*J*_ matrix) by sampling from a Bernoulli distribution for each element in the **P**_*K*×*J*_ matrix.

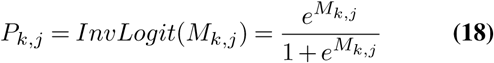

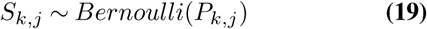

We randomly drew a set *A* = {*A*_1_, *A*_2_, *A*_3_, *A*_4_, *A*_5_} of 5 active regulatory proteins (with non-zero effects on changes in the inclusion rates) from the whole set of regulatory proteins *R* = {1, 2, …, *J*}. The regulatory effect for RBP *j θ*_*j*_ was then drawn from a uniform distribution between -2.5 and 2.5 if *j* belonged to the set of active regulatory elements *A* and set to zero otherwise. These values of *θ*_*j*_ represent the mean increase in the log(odds ratio) of exons having a binding site for that protein compared with those without a binding site.

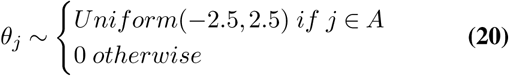

Once the parent nodes of the DAG were simulated, we could easily simulate the final data by sampling parameter values along the graph according to our model. First, we drew changes in the logit-transformed inclusion rates *β*_*k*_ from a normal distribution with mean obtained from a linear combination of effects 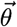 and binding sites 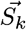 and standard deviation *ν* = 0.1. This way we introduced noise with small random changes in inclusion rates of exons that were not targets of any of the differentially active RBP.

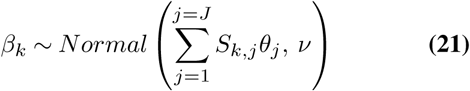

We then combined *α*_*k*_ and *β*_*k*_ to obtain the mean *logit*(Ψ) for condition *b*, and sample 3 samples from each mean using *σ* = 0.2 to introduce some inter-individual variability. Being *D*_*i*_ a variable that takes value 1 when sample *i* belongs to condition *b* and 0 otherwise,

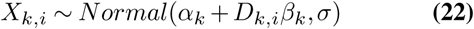

The total number of reads mapping to each event *T*_*k,i*_ were drawn from a Poisson distribution with *log*(*λ*) = 2 by default,

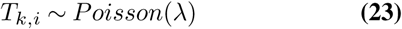

They were subsequently used to sample the corresponding reads supporting inclusion *I*_*k,i*_ from the binomial distribution with *p* = Ψ_*k,i*_, obtained from the inverse logit transformation of *X*_*k,i*_.

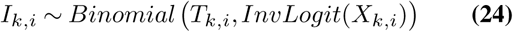

Using these default parameter values, we additionally simulated data for increasing sequencing depths (from *log*(*λ*) = 1 to *log*(*λ*) = 5.5) and with an increasing number of total regulatory proteins (from J=50 to J=250), maintaining a total of 5 differentially active RBPs to evaluate the effect of this variables on the methods performance.

### Bayesian inference

The probabilistic models were implemented in Stan (10) using non-centered parametrization, whenever it was possible, to improve sampling efficiency (7). The joint posterior distributions of the parameters were approximated using No-U Turn Sampler (NUTS) as implemented in Stan (20), running 4 chains along 4000 iterations, being 2000 of them for warming up. Convergence of the Markov Chain Monte Carlo (MCMC) algorithm was checked in each case by means of the split Gelman-Rubin R 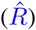 (16).

### Differential splicing analysis

In order to identify exons with significant changes in inclusion rates, a GLM with binomial likelihood was used to model the probability of inclusion of a particular exon using the sample condition *D*_*i*_ as only predictor. After fitting the model, we extracted the estimate and p-value for the coefficient representing the condition of interest. We then obtained adjusted p-values by means of Benjamini-Hochberg (BH) multiple test correction.

#### Over-representation Analysis (ORA)

We tested over-representation of binding sites for a particular RBP on the set of significantly changed exons using a Generalized Linear Model (GLM) with binomial likelihood to model the probability of being significantly changed as a function of the presence of a binding site for a particular RBP. We then extracted the p-value for the coefficient for each RBP and applied BH multiple test correction.

#### Gene Set Enrichment Analysis (GSEA)

We implemented an in-house algorithm for GSEA in python following (50). We sorted exons according to the estimated coefficient representing log-transformation of change in exon inclusion odds between the two conditions under study. We then used the matrix with binding sites for each exon and RBP and subtracted the mean for each column. This way, we give weight to each binding site depending on the number of binding sites present for a particular RBP. We then calculated the cumulative sum and took the maximum and minimum values as enrichment scores. We permuted 10000 times the list of exons to calculate a null distribution of enrichment scores, estimated p-values as the proportion of permutations with bigger enrichment scores and performed BH multiple test correction.

### Regulatory features: CLiP-seq derived RBPs binding sites

CLiP-seq binding sites were collected from several databases and merged in a single BED file (8, 14, 30, 57). Human binding sites and mouse binding sites in mm10 were transformed to mm9 coordinates using liftOver tool for compatibility with vast-tools. For simplicity, only binding sites mapping to the 250bp upstream or downstream the alternative exons were included in the analyses.

### Bench-marking of differential splicing methods using real data

In order to assess the performance of dSreg in real biological data, we used the GSE112037 dataset, which contained an independent quantification of exon inclusion rates using RASL-seq for the quantitative evaluation of the performance of different methods (58). We also evaluated the impact of sequencing depth on the performance of the different methods by serial down-sampling of sequencing up to 1/512 times the original depths (120M reads). dSreg was run using processed event counts as provided by DARTS, which is itself based on rMATS (47, 58). GLM analysis was also performed using the same event counts. MISO and BRIE were run using their own event annotation, corresponding to hg19 genome version and Ensembl annotation release 75 for all methods (21, 24). An additional *Nullmodel* for dSreg without regulatory information, as in the simulations, was run to test the improvement in detection of splicing changes by including regulatory features. For evaluation, we selected events with at least 50 total reads in the RASL-seq experiment, and calculated the real inclusion rates as the proportion of reads supporting exon inclusion. Real AS changes were defined as those with a |ΔΨ| > 0.05 and *FDR* < 0.05 using a basic GLM in R. Then, performance was evaluated by comparing the estimation of the ΔΨ in the down-sampled RNA-seq experiments and the ones derived from RASL-seq. We assessed the quantitative estimation of inclusion rates by calculating the Pearson coefficient with the real ΔΨ. AUROC was used to asses the ability to identify differentially spliced. The scoring function for AUROC calculation were: i) Bayes Factors for BRIE and MISO: 1 - FDR for GLM; ii) and *P* (|ΔΨ| > 0.05|*data*) for DARTS and dSreg; and iii) MISO and BRIE were evaluated using only the subset of events that were also represented in the RASL-seq experiments.

### Assessment of the ability of dSreg to identify AS regulatory drivers using ENCODE knock-down experiments

In order to evaluate the performance of dSreg in detecting the RBPs that drive AS changes between two conditions, we used the data from systematic knock-down experiments of 206 RBPs in two different human cell lines from the ENCODE project and their corresponding binding profiles (14, 37). We downloaded the rMATS processed files available from the website and analyzed their regulatory patterns using GLM+ORA, GLM+GSEA and dSreg. Regulators were defined as differentially active if FDR<0.05 for both ORA and GSEA; or if the posterior probability of the *θ*_*j*_ being different from 0 was higher than 95% (*P* (|*θ*_*j*_| > 0|*data*) > 0.95) for dSreg. The performance was evaluated with 3 different measures. First, we analyzed the number of times the RBP that was down-regulated was found among the driver regulators of AS. This measure was normalized by the expected proportion of matches if the regulatory elements were selected randomly from the set of available regulators. Second, we measured the proportion of RBPs defined as differentially active were differentially expressed in in the knock-down experiment. Third, we sorted by absolute differential activity (or FDR for ORA and GSEA) the RBPs and calculated of RBP that was knocked-down.

### Real data analysis

GSE59383 fastq data were downloaded and mapped using vast-tools 0.2.0 (23) to identify AS events. We restricted our analysis to exon cassette events that showed at least 1 inclusion and skipping read in at least one sample. Once extracted the number of inclusion and total counts for each event and sample, we used all the methods described here (ORA, GSEA and dSreg) to analyze regulatory patterns using a compendium of CLiP-seq binding sites.

## Results

### Adding information about regulatory elements improves the detection of AS changes even at low sequencing depth

Using simulated data we first compared the performance of a standard GLM to detect changes in AS at different sequencing depths (*λ*). Correlation between the estimated 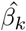 and the real *β*_*k*_ used for the simulations was generally low and did not increase with sequencing depth. When focusing on detection of AS changes, we found that, as expected, at low sequencing depths (*log*(*λ*) ≤ 3), the sensitivity at a 5% False discovery rate (FDR) was smaller than 10% when using a simple GLM. As *λ* increased, so did the sensitivity of the GLM (Fig. 2B). Interestingly, the F1 score, which integrates both sensitivity and specificity, saturated with depth, suggesting that after some point, there was not much gain by increasing sequencing depth (Fig. 2C). To avoid the need to select an arbitrary threshold to assess the performance of the different methods, we additionally calculated the Receiver Operating Characteristic (ROC) curves for each simulated dataset and the area under them (AUROC, Fig. 2D and E). These results showed that, at low sequencing depths (*log*(*λ*) < 3), the performance was rather poor, with AUROC values of 0.7 at most.

**Fig. 2.**
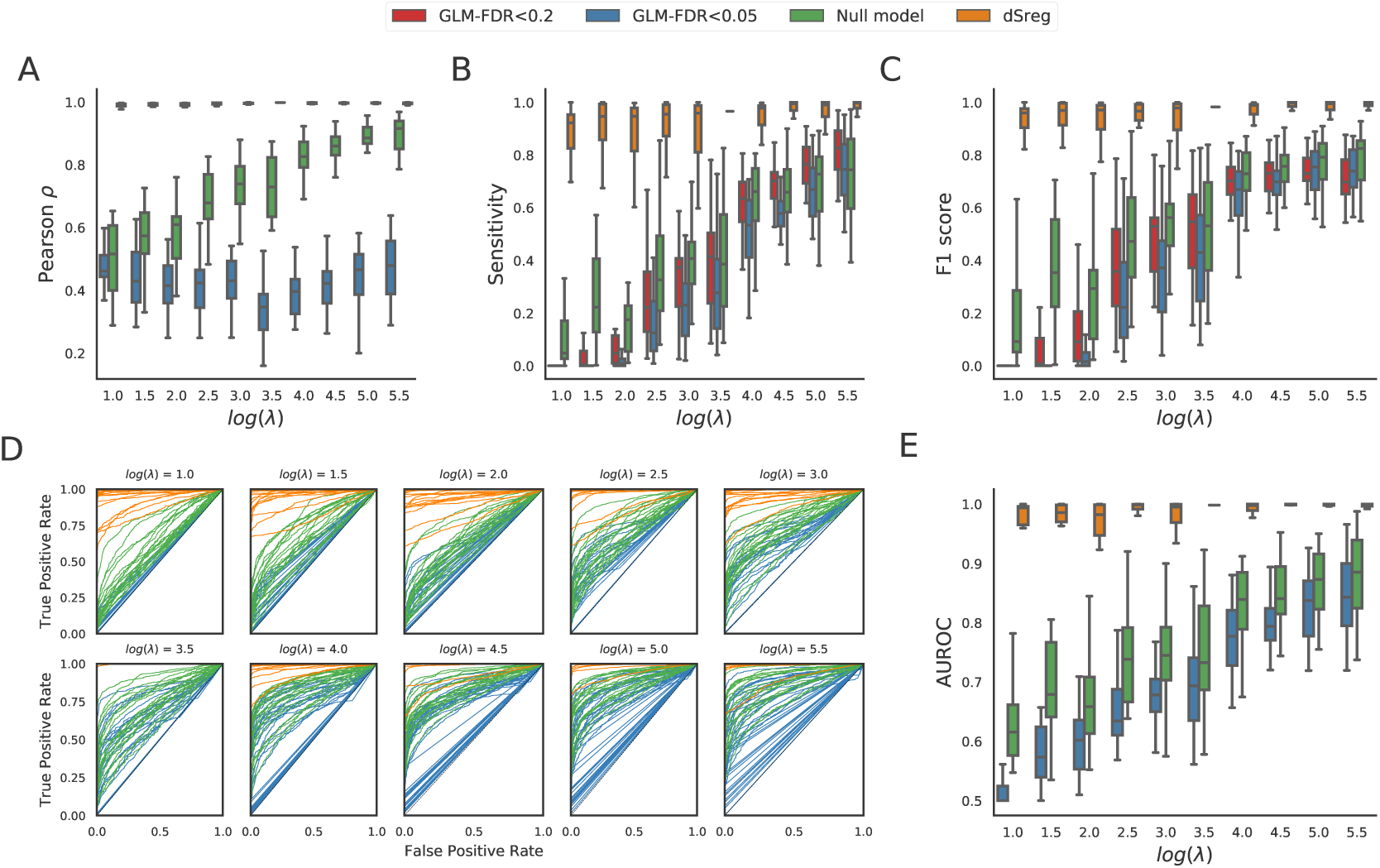
Comparison of the performance for the identification of different event inclusion rates of a standard method using a single GLM per exon considering two FDR thresholds (0.05 and 0.2),a bayesian model that pools variance across all exons (*Null model*) and dSreg. Performance was analyzed in simulations with increasing sequencing depths *λ* (the mean of the Poisson distribution used to simulate the total number of reads mapping to an exon skipping event). **A**. Pearson correlation between real and estimated *β*_*j*_. **B** Sensitivity. **C** F1 score. **D, E** Receiver Operating Characteris.tic (ROC) curves (D) and the area under them (E)

In order to check whether potential improvements of dSreg were due to the inclusion of binding sites and changes in RBPs activity in the model or just to variance pooling, we ran dSreg and a reduced model that only pools variance from all exons without taking into account of the binding sites and changes in regulatory activities (*Null model*). We defined as significantly changed events as those with a posterior probability higher than 95% of having a *β*_*k*_ > 0. The *Null model* already outperformed the GLM at the single exon level and improved quantitative estimation of *β*_*j*_ with depth (Fig 2A). However, dSreg showed a much greater improvement in correlation and sensitivity, even at very low sequencing depths (*log*(*λ*) < 3), when there was practically no information from individual events (Fig. 2). This increased sensitivity did not come with a decrease in specificity as could be expected, since it showed also very high F1 scores and AUROC, suggesting that differences in performance are intrinsic to the method and not threshold dependent (Fig. 2C,D and E). Results with the *Null model* suggest that pooling variance across events does only marginally improve the inference of splicing changes, at least with the low variance used in these simulations. dSreg, in contrast, additionally used the information about the underlying regulatory mechanisms to correct differences that may easily arise by chance in datasets with limited sample size, given that simulations were done with only 3 samples per condition.

### dSreg improves the detection of the RBPs driving AS changes

Once AS changes have been identified, we focused on the detection of the regulatory elements potentially controlling these events. Using our simulated datasets, we compared dSreg with the traditional ORA and GSEA approaches. As FDR<0.2 filtering showed higher F1 score in the identification of splicing changes (Fig. 2C), we used this threshold to select significantly changed events to perform the downstream enrichment analyses. The dependency of ORA on the detection of significant changes led to low F1 scores for GLM results at any tested FDR threshold, especially at low sequencing depths (Fig. 3A). We also used an in-house version of GSEA to take advantage of quantitative information in the identification of regulatory elements. Briefly, events were ranked according to their Maximum Likelihood Estimation (MLE) of the coefficient of the GLM, which represents the log of the odds ratio of inclusion between the two conditions. Then, we looked for non-random distributions of binding sites along the ranked list (50) (see Methods section for details). We found a substantial improvement over ORA, with higher F1 scores, especially at low sequencing depths, but did not seem to benefit from higher sequencing depths (Fig. 3A). dSreg outperformed both ORA and GSEA at every evaluation metric, and was barely affected by low sequencing depths (Fig. 3). Therefore, integration of the two sources of information improves results both in terms of inference of differential inclusion rates and the identification of the mechanisms driving those changes.

**Fig. 3.**
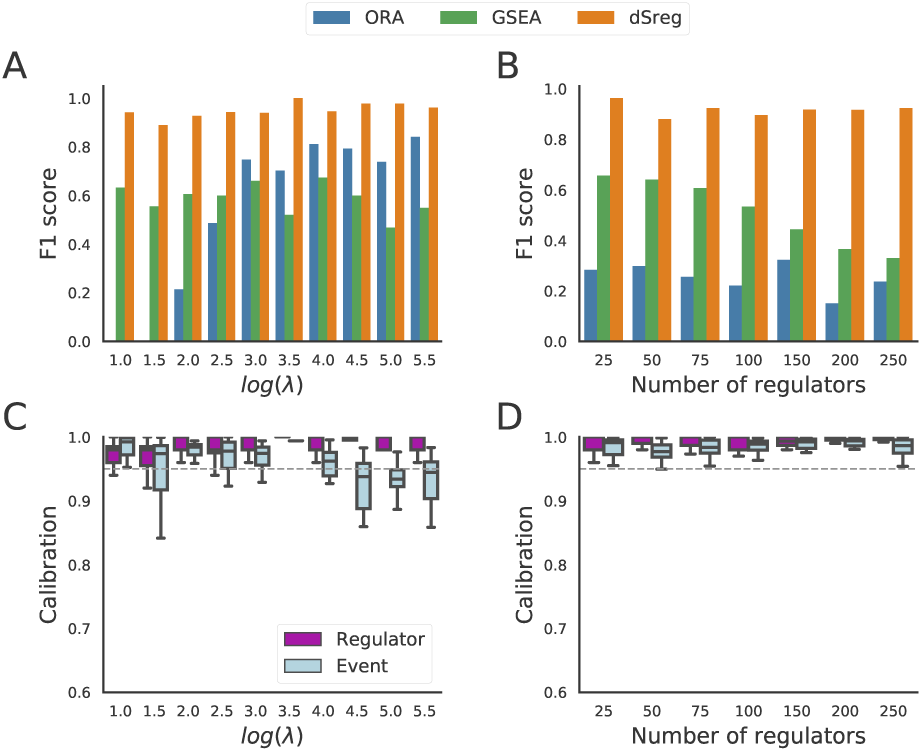
Performance of methods for the detection of regulatory elements: ORA with variable FDR thresholds (0.05 and 0.2), non-parametric GSEA and dSreg. Performance was analyzed in simulations with increasing sequencing depths *λ*, which is the mean of the Poisson distribution used to simulate the total number of reads mapping to an exon skipping event. **A, B**. Mean F1 scores obtained with different depths *λ* (A) and total number of regulators (B) for the different enrichment approaches. **C, D** Calibration, measured as the proportion of times the real value lies within the 95%CI of differentially spliced exons and regulatory elements for increasing sequencing depth (C) or increasing number of total regulatory elements (D).

### Increasing the number of potential regulatory elements does not decrease dSreg performance

We have so far used simulated data to explore the effect of sequencing depth on both the detection of splicing changes and on the identification of the key RBPs driving these changes. We next assessed the impact of the number of regulators, which may increase the number of false positives, particularly in presence of co-linearities among binding profiles of different RBPs. To study this potential limitation, we simulated datasets with only 5 active RBPs as in the previous simulations, but increasing the number of total RBPs included in the analysis up to 250. We found that the F1 score tended to decrease as the number of potential regulators increased with either ORA or GSEA, despite multiple test correction to control false discovery rate. Once more, dSreg outperformed both methods and remained unaffected by the inclusion of other inactive regulatory elements (Fig. 3B).

### Model calibration remains robust while decreasing the proportions of active RBP

We further analyzed the performance of dSreg in terms of calibration. A model is well calibrated when inferred probabilities actually represent the real frequency of a given phenomena i.e. a model is calibrated when the uncertainty of the parameter estimate matches the evidence contained in the data. Calibration was calculated as the proportion of events and regulators whose real change in logit-transformed inclusion rates (*β*_*k*_) or activity (*θ*_*j*_) is within the estimated 95%CI. Whereas changes in inclusion rates were well calibrated, the uncertainty of the changes in the activity of RBPs seemed to be slightly overestimated, given that 95%CI included the real values more often than 95% of the times, independently on the sequencing depth *λ* (Fig. 3C). We then tested how different numbers of total regulatory elements affected model calibration with the previous simulations using only 5 active out of an increasing number of candidate RBPs. We found that the total number of candidate regulators had no effect on calibration (Fig. 3D). These results suggest that dSreg is conservative when estimating the uncertainty of the regulatory activities *θ*_*j*_ based on the data, since the real value is within the 95%CI more often than expected across all tested conditions (Fig. 3C,D).

### dSreg outperforms other methods using real data

To assess whether the better performance of dSreg could be confirmed with independent real data, we used an RNA-seq dataset (around 120M reads per sample) for which a subset of AS events were quantified using RASL-seq and can be used as gold standard (58). We used CLiP-seq data of a number of RBPs binding to upstream and downstream flanks of exon skipping events as regulatory features for dSreg (14). Since dSreg performed particularly better than other methods at low sequencing depths, we subsampled the sequencing reads by a factor of 2 up to 512 to analyze the extent of this advantage. We analyzed the data also with MISO, BRIE and DARTS. Both BRIE and DARTS use prior information to improve detection of splicing changes (21, 24, 58). dSreg and the *Null model* showed the best performance, compared to all other methods, except in extremely low coverages (dilution factor > 100), in which DARTS overcame dSreg (Figure 4A,B). In contrast to the results obtained from the simulated data, dSreg and *Null model* performed similarly, which suggests that the regulatory features that were added do not contribute much to the estimation of AS changes. However, it also shows that it remains robust to the inclusion of non-relevant regulatory features. Neither BRIE nor DARTS out-performed the *Null model*. We observed the same patterns when comparing the results to the full coverage RNA-seq dataset (Figure S1).

**Fig. 4.**
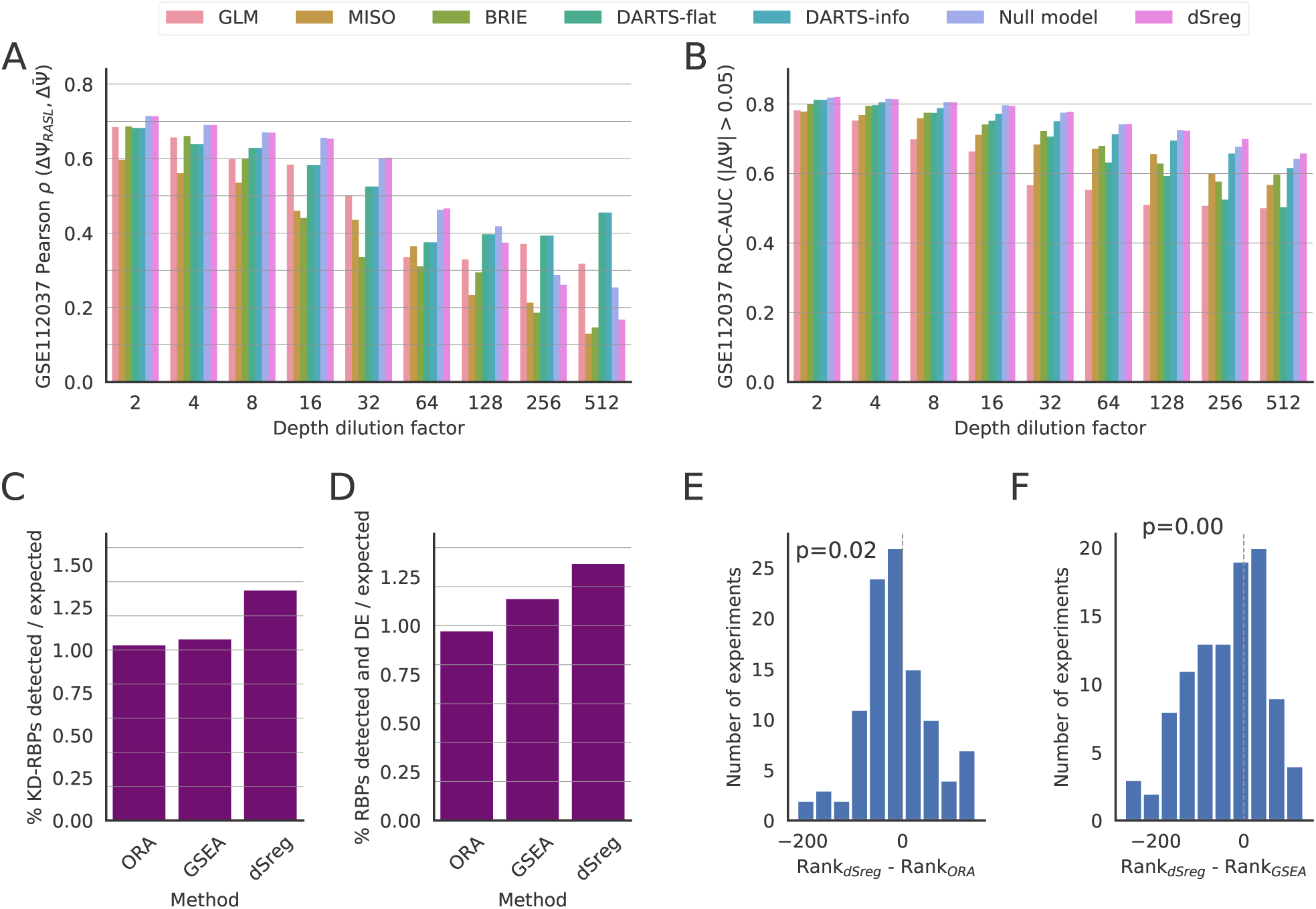
Evaluation of the performance of dSreg with other methods on real data. **A**,**B** Performance of differential splicing methods using RASL-seq quantification as true values, measured by Pearson correlation of ΔΨ (A) and Area Under the ROC (AUROC) for exons significantly changed, defined as those with a |ΔΨ| > 0.05. Methods include a GLM. MISO, BRIE, DARTS with and without using the predictions as prior (info and flat respectively), and dSreg and its *Null model*. **C** Percentage of experiments in which the knocked-down RBP was found among the regulatory elements compared to expectation. **D** Percentage of regulatory RBPs identified by each method that were detected to be differentially expressed (DE) compared to expectation. Expectations were calculated by 20000 random sampling of the same number of regulators. **E, F** Difference in rank occupied by the knocked-down RBP in the output of dSreg with that of ORA (E) and GSEA (F).

The main advantage and motivation of dSreg is the inference of the regulators driving AS changes, a feature that is not provided by any of the existing tools for AS analysis. To assess whether dSreg outperforms ORA and GSEA also with real data, we used the collection of RBP knock-down experiments from ENCODE (37). Although it is difficult to know the actual regulatory mechanisms in each case, one may reasonably assume that at least some of the AS changes would be mediated by the down-regulation of the target RBP. dSreg detected the highest percentage of knock-down RBPs as regulatory elements compared to the random expectation in each case (Figure 4C). If the expression of other regulatory element is affected by the perturbation, we would expect them also to contribute to explain AS changes. Regulators detected by dSreg tended to be more often differentially expressed in the same experiment than expected by chance compared to other methods (Figure 4D). Finally, we observed that, when sorting the regulators by their evidence, the RBP that was knocked-down tended to appear higher in the ranking produced by dSreg than in those yielded by ORA and GSEA (Figure 4E and F, respectively). Altogether, these results suggest that dSreg also outperforms previous methods in the identification of regulatory elements using real data.

### AS regulation in cardiomyocyte differentiation by core-spliceosomal factors

We then tested our model on a dataset of mouse cardiomyocyte differentiation from cardiac precursors (GSE59383) with 3 samples per condition as in our simulated scenario. Binding sites for a number of RNA binding proteins were obtained from CLiP-seq experiments and only those located in the upstream and downstream intronic flanking 250bp were used (see Extended Methods section for details). We run the 3 approaches explored in this work and found that ORA resulted in a high number of significantly enriched candidates, most of which are likely to represent false positives as in our simulation analysis (Fig. 5A). GSEA, on the other hand, showed no significant enrichment at *FDR* < 0.05, and only a few at nominal *p* < 0.05, which suggest that these p-values can easily arise by chance. Indeed, there is little concordance with results from ORA (Fig. 5A and B). dSreg did show an overall agreement with ORA results, but, as expected, dSreg provided a reduced number of RBPs whose combined action best explain the observed AS changes (Fig. 5, Table S1). Interestingly, a great deal of the identified regulatory RBPs are considered to be members of the core spliceosome (BUD13, EFTUD2, PRPF8, SF3A3, SF3B4), suggesting that changes in the activity of these particular components might be key for the AS changes underlying cardiomyocyte differentiation. In this regard, the core spliceosomal machinery has been shown to have extensive regulatory potential (54) and mutations in one of these genes (EFTUD2) have been associated with congenital heart defects, among other phenotypes (31).

**Fig. 5.**
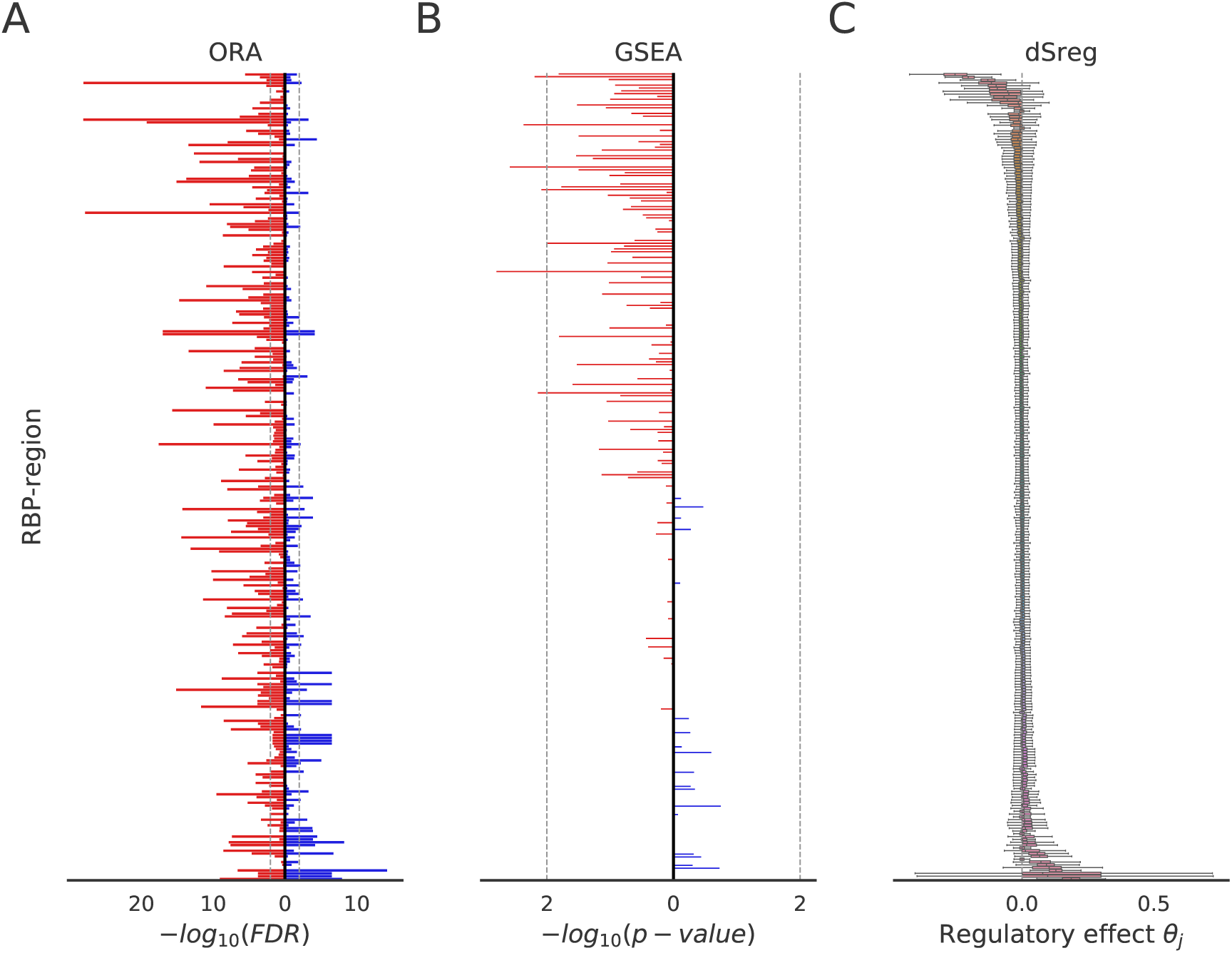
Comparison of ORA, GSEA and dSreg using a real RNA-seq dataset from a cardiomyocyte differentiation experiment. RBPs on the y-axis are sorted for the three panels according to the posterior mean of the regulatory effect *θ*_*j*_ inferred by dSreg. **A**. Candidate regulatory proteins derived from the ORA on the significantly included (blue) or skipped (red) exons represented by their significance expressed as the log transformation of the FDR. **B**. GSEA results represented by the nominal empirical p-value resulting from permuting the exon labels. RBPs with positive enrichment scores are represented on the right, and those with negative scores on the left. **C** Posterior distributions of the regulatory effects *θ*_*j*_ inferred by our model.

## Discussion

Here we present dSreg, a new method that integrates the analysis of differential AS and the identification of the underlying regulatory mechanisms in a single model. Our single-step model bypasses the need to call for differential splicing before enrichment and therefore improves sensitivity, especially at low sequencing depths. It also increases specificity as it uses information from the underlying changes in RBPs activity to avoid false positives derived from small sample sizes. Moreover, dSreg analyzes the regulatory activity all RBPs simultaneously to correct for possible co-linearities in the binding profiles and uses a horseshoe prior to force most of the RBPs activities to remain unchanged. Joint modeling also provides higher specificity in the detection of regulatory mechanisms as it reduces the number of false positives due to co-occurrence of binding sites of different RBPs, leading to an improved overall performance compared with classical enrichment approaches for regulatory elements. Our model opens the possibility to analyze AS more accurately using RNA-seq data with low sequencing depth, both for reanalysis of previously sequenced samples or for more cost-effective new RNA-seq experiments with focus on the regulatory mechanisms. Although transcript-based methods also lower the requirements on sequencing depths (1, 53), our model works directly at the event level, reducing the dependency on the transcript annotation (59). In contrast to previous approaches, including bayesian methods like MISO (24), our model is motivated by how splicing changes are regulated between two biological conditions rather than on how inclusion and skipping reads are generated from the inclusion rate (Ψ) in a particular sample. Still, we show that not only we gain more biological insight directly from the model, but also obtain, at least, as good estimations of AS changes as provided by the best performing tools to date. dSreg still requires a previous definition of alternative mapping events and allocation of reads to inclusion or skipping isoforms, making it compatible with any of the software used in this paper.

Our good results on simulations are, however, restricted to those cases in which AS changes are mediated only by a subset of differentially active RBPs binding to known sites. Although inclusion of a high number of RBPs showed no effect on the changes in inclusion rates between the two tested conditions, alternative sources of errors, such as errors in the binding profiles or missing information might have a negative impact on the sensitivity of dSreg. Indeed, the improvement of dSreg on real data compared with the *Null model* is rather small, if any. The better performance of both models compared with existing methods seems therefore due to the inclusion of a parameter describing variance between samples across exons rather than to the regulatory information. Similarly, other methods including event features to improve detection of splicing changes do not outperform our *Null model* (21, 58), except when data is very scarce. Whereas dSreg only uses AS data from the target experiment, DARTS informative prior was trained with many other datasets such that, in absence of information, is able to make relatively good predictions about the outcome of an experiment given the regulatory features. These results suggest that only a small part of splicing variation is mediated by RBPs CLiP-seq binding sites as in the model. This was not unexpected, since previous studies suggest that AS regulation is far more complex than a sum of effects of a number of RBPs and that RNA structure plays a critical role (6, 29, 51). In spite of these limitations, dSreg was able to detect the knocked down and differentially expressed RBPs in loss of function models more efficiently than traditional approaches like ORA and GSEA, suggesting that the identified regulation was real. Yet, we expect that careful modeling of additional AS regulatory features will improve the results, e.g. nucleosome positioning and histone modifications (22, 32, 34). Moreover, dSreg is limited so far to pairwise comparisons, whereas we are often interested in analyzing enrichment over a number of conditions, such as time series and dose-response experiments. Further work will be necessary to allow for more complex and powerful experimental designs.

## Conclusion

Our model provides an example of how joint modeling of interdependent phenomena can improve results compared with completely separated analysis relying on categorization according to rather arbitrary thresholds. Bayesian inference through MCMC methods provides a general framework to fit very flexible models that adapt to each particular analysis and to easily extend currently existing models to integrate different sources of information. In our case, we only integrated binding sites information with AS data, but these models are flexible enough to include information about expression of AS regulators, post-transcriptional modifications or any other piece of information supporting a change in the activity of a particular regulatory protein. This model is not only limited to regulation analysis, but can also be used with functional annotations, such as the presence of functional domains, phosphorylation sites, protein-protein interaction motifs, or any other property that may be associated with AS. Moreover, we have implemented the model in dSreg (https://bitbucket.org/cmartiga/pydsreg/src/master/), which enables running the model using only the matrices of inclusion and total number of reads per event and a matrix *S* with the event features e.g. the binding sites. Therefore, dSreg adds a valuable statistical tool to existing software aimed at identifying AS events, such as rMATS or vast-tools (23, 46), among others, for more accurate detection of AS regulatory mechanisms using RNA-seq data.

## ACKNOWLEDGEMENTS

We would like to thank Victor Jimenez for critical reading and useful discussions about the manuscript and beyond. We also thank Eric Van Nostrand for his help to retrieve the processed ENCODE data.

