## Supplementary figures and images for "dSreg: A bayesian model to integrate changes in splicing and RNA binding protein activity"

### Figure S1

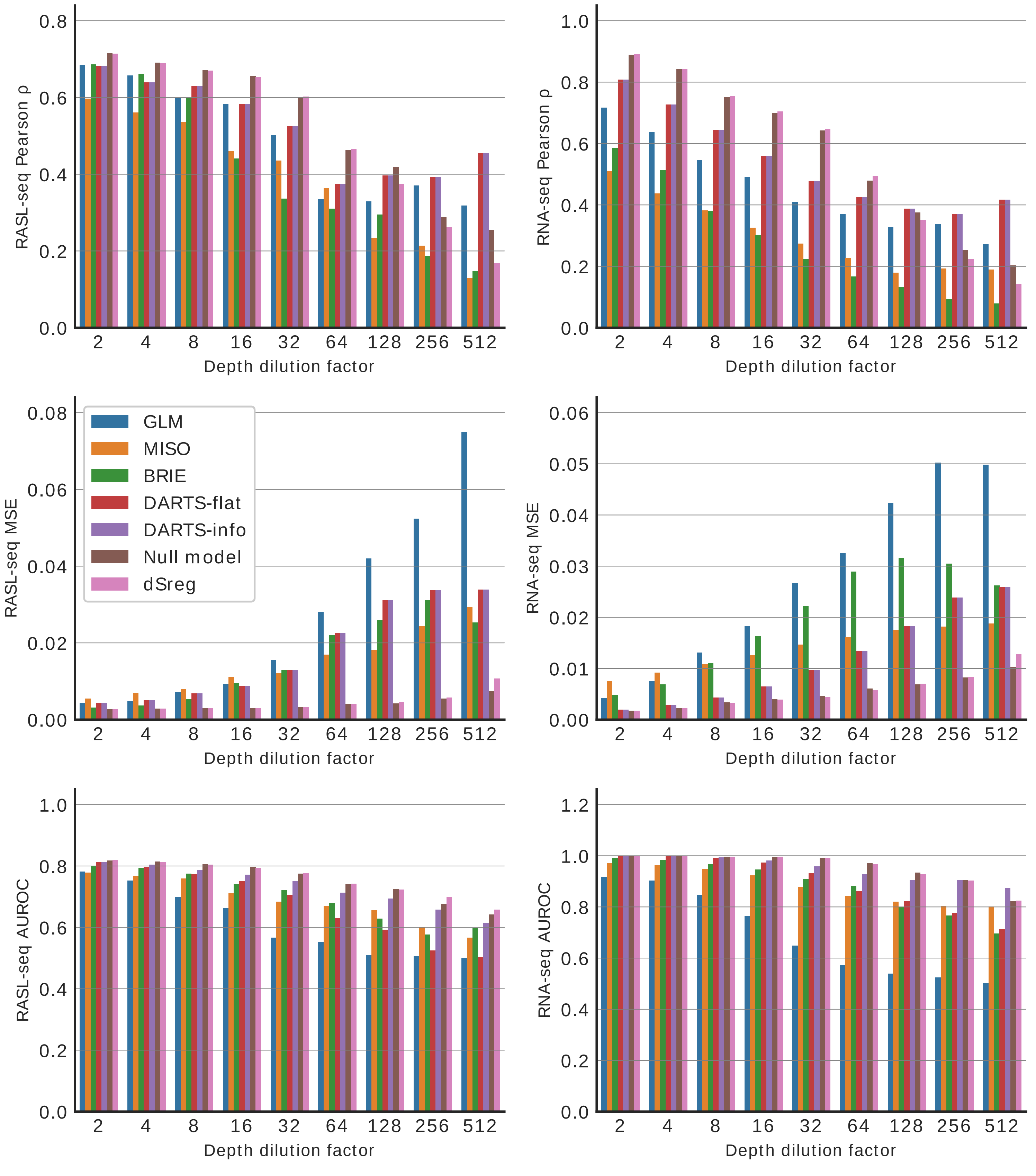
